# Scarless regeneration requires post-translational proteolytic regulation via macrophage-derived Legumain

**DOI:** 10.64898/2026.10.20.760451

**Authors:** Julia Mo, Hailey Levitz, Safiyah Ali, Subin Lee, Desmarie Sherwood, Troy Anderson, Katherine Spack, Maria Blumenkrantz, Everett Weinstein, Joanna Smeeton

## Abstract

Macrophages express proteolytic factors in response to injury but how proteases are deployed and regulated during regeneration are incompletely defined. Beyond their transcriptomic expression, activity of individual proteases is defined both by their unique substrate specificity and post-translational modifications of their active forms. Here we show a key role for the lysosomal protease Legumain in controlling ECM dynamics and post-translational processing of the regenerative proteome. Injury interface-targeted Ctsla localization and post-translational processing requires Lgmn in macrophages. Extracellular cues from physical remodeling control differentiation state and injury response of ligament resident fibroblasts. Loss of ECM turnover, including localized Fibronectin 1a/b remodeling, impairs dedifferentiation in legumain mutants and leads to fibrotic scarring. Taken together, we demonstrate a role for macrophage-derived lysosomal proteases as specialized tools required to fine-tune proteome composition and function in regeneration.

## Introduction

Regeneration is a complex process requiring the coordination of multiple cellular events to occur in tandem or sequentially throughout the course of repair. Adult zebrafish have a high capacity for endogenous regeneration, demonstrated by the growing number of injury models showcasing their regenerative capacity in the heart, spinal cord, caudal fin, and jaw joint^1–6^. ECM-rich connective tissues such as ligaments and tendons are challenging to rebuild, often repairing with inferior scar tissue, increasing the susceptibility of damage-associated joints to osteoarthritis^7–9^. Our current understanding of the underlying biology of regeneration largely lies at the epigenetic and transcriptomic level, yet there is a gap between gene expression and proteomic function that must be addressed. Post-translational modifications (PTMs) serve as key regulatory mechanisms that expand proteomic diversity and govern cellular function. Here, we show post-translational protein regulation of the injury microenvironment is mediated through macrophage-derived proteases. Using a *legumain* mutant model of fibrotic healing, we uncover how proteases shape tissue architecture to support scar-free ligament regeneration.

Immune contributions to the ECM are essential for healing. In response to injury, a temporary matrix is assembled to bridge the injury site and serve as a scaffold for immune cells to migrate throughout the wound. Fibroblasts are the main source of matrix deposition and macrophages are best known to coordinate ECM remodeling through crosstalk with fibroblasts^10–13^. Our knowledge of macrophage-mediated ECM remodeling have expanded with recent work showing that zebrafish and mouse macrophages can directly contribute collagen and fibronectin in response to heart and ligament injury^7,14,15^. Additionally, Collagen V deposition by macrophages is essential for collagen assembly to initiate scar formation after myocardial injury in mice^15^. Furthermore, macrophage-mediated ECM remodeling can influence cell behavior. This influential role has been observed in zebrafish after myocardial cryoinjury where *mmp14b+* macrophages home to the injury border and promote cardiomyocyte invasion through ECM degradation^2^. ECM deposition by immune cells has also been observed from neutrophils after bone fracture, where they secrete insoluble cellular fibronectin to initiate granulation tissue formation in the fracture gap^16,17^. Neutrophils and macrophages are the first responders to injury, often producing high levels of lysosomal proteases to facilitate critical processes such as adaptive immune cell activation, ECM remodeling, cell migration, and apoptosis^18,19^. Proteolytic processing is instrumental for many biological processes including protein turnover, antigen presentation, cell migration, apoptosis, and ECM remodeling^20–28^. Legumain (Lgmn) is a cysteine endopeptidase with a unique sequence-specificity for asparagine residues^29^. Its ability to release, activate, or degrade specific protein targets, as well as its pH-dependent auto-activation, designate it as a specialized molecular tool^29–31^. A mechanistic understanding of how substrate-specificity of different proteases are exploited to precisely reshape the proteome and influence tissue topography remains poorly defined.

We have previously shown a requirement for Lgmn and ECM remodeling for scarless ligament regeneration^7^. After ligament injury, newly synthesized tissue is highly disorganized in *legumain* mutant zebrafish (*lgmn^MUT^*), and they heal with a dysmorphic scar^7^. Further, we observed *lgmn+* and *fn1a+* macrophages wrapped around the injury edge of the ligament^7^. Ligament repair provides a complex tissue topography to investigate the mechanisms involved in reshaping the terrain, such that the dense mature ECM is degraded while also protecting the new ECM. We hypothesized that Lgmn-mediated ligament breakdown is essential to establish an interface between old and new matrix for their integration into a unified scaffold used for subsequent tissue patterning in regenerative healing.

In this study, we dissected the mechanisms underlying Lgmn-mediated tissue remodeling during ligament regeneration. Using scRNAseq transcriptomic and LC/MS proteomic approaches, we uncovered a role for Lgmn in reshaping the proteome through precise and targeted proteolytic regulation of Fibronectin 1a/b (Fn1a/b) and Cathepsin La (Ctsla). Although global protease abundance was decreased in *lgmn^MUT^* following injury, there was no functional impairment of global protease activity. We determined that Ctsla abundance is elevated to compensate for Lgmn-deficiency. Although increased levels of Ctsla is sufficient to rescue global protease activity, Ctsla may be proteolytically degrading with less specificity due to its less stringent requirements for proteolysis compared to Lgmn^30,32^. Moreover, we show that Lgmn is involved in processing Ctsla from its single-chain to double-chain form, which can further modify protease activity and substrate specificity. Together, this work demonstrates how lysosomal proteases are deployed during regeneration as specialized tools required to fine-tune proteome composition and tissue integration.

## Results

### Lgmn abundance peaks during early healing

To determine the temporal dynamics of active Lgmn during ligament regeneration, we transected the interopercular-mandibular (IOM) ligament connecting the interopercular bone (IOP) to the retroarticular process (RA) of the lower jaw joint in adult zebrafish (Figure 1a). We used western blot analysis to determine when Lgmn was most abundant after injury by generating protein lysates from whole-resected joints collected from uninjured, 6 hour, 1, 3, and 7 days post ligament transection (dplt) in WT and *lgmn^MUT^*. Using an antibody against Lgmn, WT samples had bands present at both 36 and 25 kDa, which are the expected sizes of activated Lgmn (Figure 1b). As expected, no bands were detected in any of the *lgmn^MUT^*samples, confirming the mutation encoding for a premature stop codon results in a loss of Lgmn translation (Figure 1b). These data showed that in WT, Lgmn levels significantly increase at 1 and 3 dplt compared to uninjured levels (p=0.0048 and p=0.0003, respectively) and wane by 7 dplt (p=0.5204) (Figure 1c). Taken together, this temporal expression pattern indicates Lgmn function most likely occurs within the first 3 days after injury.

**Figure 1.**
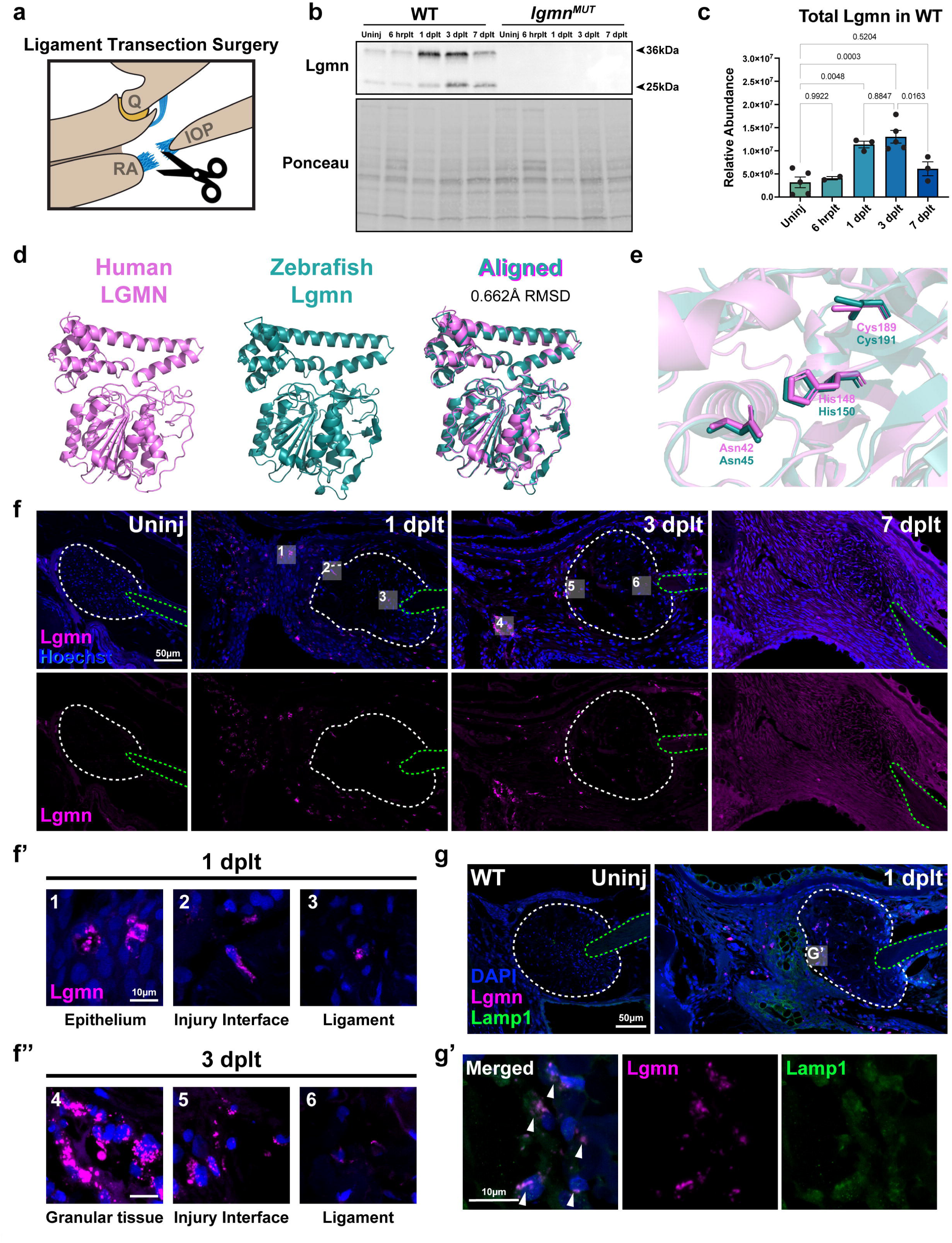
Active Legumain is present after injury. a) Schematic of unilateral IOM ligament transection surgery in adult zebrafish. RA = Retroarticular, Q = Quadrate, IOP = Interopercular Bone. b) Western blot of 36kDa and 26kDa active forms of Lgmn in lysates generated from whole-resected joints collected from WT and *lgmn^MUT^* at uninjured, 6 hrplt, 1, 3, and 7 dplt timepoints. c) Quantification of Lgmn western blot from (b), relative intensity of both bands were measured (n=2-5 per timepoint). Error bars = SD. d) 3D structures of human proLGMN (magenta), zebrafish proLgmn (teal), and cross-species alignment of both structures. RMSD = Root Mean Squared Deviation. e) Pairwise alignment of human LGMN (magenta) and zebrafish Lgmn (teal) showing the residues of the catalytic triad. Asn=Asparagine, His = Histidine, Cys = Cysteine. f) IF of Lgmn (magenta) counterstained with Hoechst (blue) in WT uninjured, 1, 3, and 7 dplt samples (n=3-8 per timepoint). (f’, f’’) High magnification images of Lgmn localization at 1 dplt (f’) and 3 dplt (f’’) from areas marked by transparent white boxes 1-6 from (f). White dashed-lines marks ligament. Green dashed-line marks IOP bone. g) Colocalization of Lgmn and Lamp1 visualized through multiplex of Lgmn-tsa (magenta) and Lamp1 IF (green) counterstained with DAPI (blue). White dashed-line marks ligament. (g”) Inset marked in (g), arrows mark co-localization. Scale bars = 50µm (f, g), 10um (f’,f’’, g’).

### Structural alignment of Zebrafish Lgmn to Human LGMN determine conservation

Previous work has shown that both the 36kDa and 25kDa activated forms of human LGMN harbor enzymatic activity^33^. Lysate from zebrafish joints have bands of the same sizes, suggesting a conservation in Lgmn activation between the two orthologs. We next sought to determine the level of evolutionary conservation of both the overall structure and functional core of zebrafish Lgmn (zLgmn) compared to human LGMN (hLGMN) (Figure 1d-e). Analysis of amino acid identity between zLgmn and hLGMN were 65% similar. Pairwise structural alignment of 3D crystal structures between the alpha-fold predicted zLgmn and experimentally derived hLGMN structure shows high conservation, with a root mean squared deviation (RMSD) value of 0.662Å (Figure 1d). A RSMD value <2.0Å suggests a high degree of similarity due to highly aligned protein structures, therefore there is high conservation of structure between species. Moreover, structural and sequence alignment of active site residues demonstrated that the catalytic triad residues of zLgmn (Asn45, His150, Cys191) are positionally equivalent to hLGMN (Asn42, His148, Cys189) (Figure 1e). These alignments suggest conservation of structure and functional site between mammalian LGMN and teleost zebrafish Lgmn. Therefore, catalytic function is likely conserved.

### Lgmn levels are broadly elevated throughout the injury site

Since structural analysis determined that Lgmn is likely functioning as a protease in the context of ligament regeneration, we sought to identify the spatial distribution of Lgmn within the injury site. Through immunofluorescence (IF), we observed that Lgmn is present throughout the regenerate tissue. Specifically, we observed localization within the skin domain, granular tissue, and the injured ligament at 1 and 3 dplt, with no Lgmn detectable in those domains by 7 dplt (Figure 1f, f’, f”). Lgmn+ cells are present surrounding the ligament injury border as well as within the injured ligament at 1 and 3 dplt (Figure 1f’, f”). These sites of concentrated Lgmn suggest that a subset of Lgmn+ cells directly target the injured tissue for Lgmn-mediated proteolysis. As Lgmn is canonically a lysosomal protease in mammals, we confirmed if this subcellular localization was comparable in zebrafish. Using multiplexed tyramide signal amplification (tsa) of Lgmn with Lamp1 IF to mark lysosomes, we observed co-localization of Lgmn and in Lamp1+ lysosomes, including at the injury interface (Figure 1g, g’). Taken together, these data show that Lgmn is found in the acidic environment that facilitates Lgmn autoactivation and suggest that Lgmn has a lysosomal function during ligament regeneration.

### *lgmn^MUT^* have a pro-fibrotic transcriptional response to ligament injury

Next, we tested whether *lgmn^MUT^* fail to regenerate due to the presence of an aberrant cell population or an impaired transcriptional response to injury. We used single-cell RNA sequencing (scRNAseq) to identify the cellular populations present and profile the transcriptional response. scRNAseq datasets were generated by collecting whole joints from WT and *lgmn^MUT^* at uninjured, 1, 3, and 7 dplt. The tissue was dissociated into single-cell suspensions, and live cells were selected by FACS and input onto the 10x Chromium platform for sequencing (Figure 2a). After processing and QC filtering, we recovered 18,079 cells (7,635 WT, 10,444 *lgmn^MUT^*) with a median of 2,166 genes and a median of 14,707 reads per cell (Table S1). From low resolution clustering of the data using Seurat^34^, we identified 2 endothelial (*acta2a+, col5a3a+, pecam1a+, kdrl+)*, 11 epithelial, 9 skeletal, and 7 immune-related (*mpx+, lyz+, mpeg1.1+, spl1b+, gata2a+, cd37+, gata3+*) clusters (Figure 2b, S1b). No clusters were exclusive to one genotype, indicating that *lgmn^MUT^*had similar cellular composition compared to WT at the same timepoint (Figure S1a).

**Figure 2.**
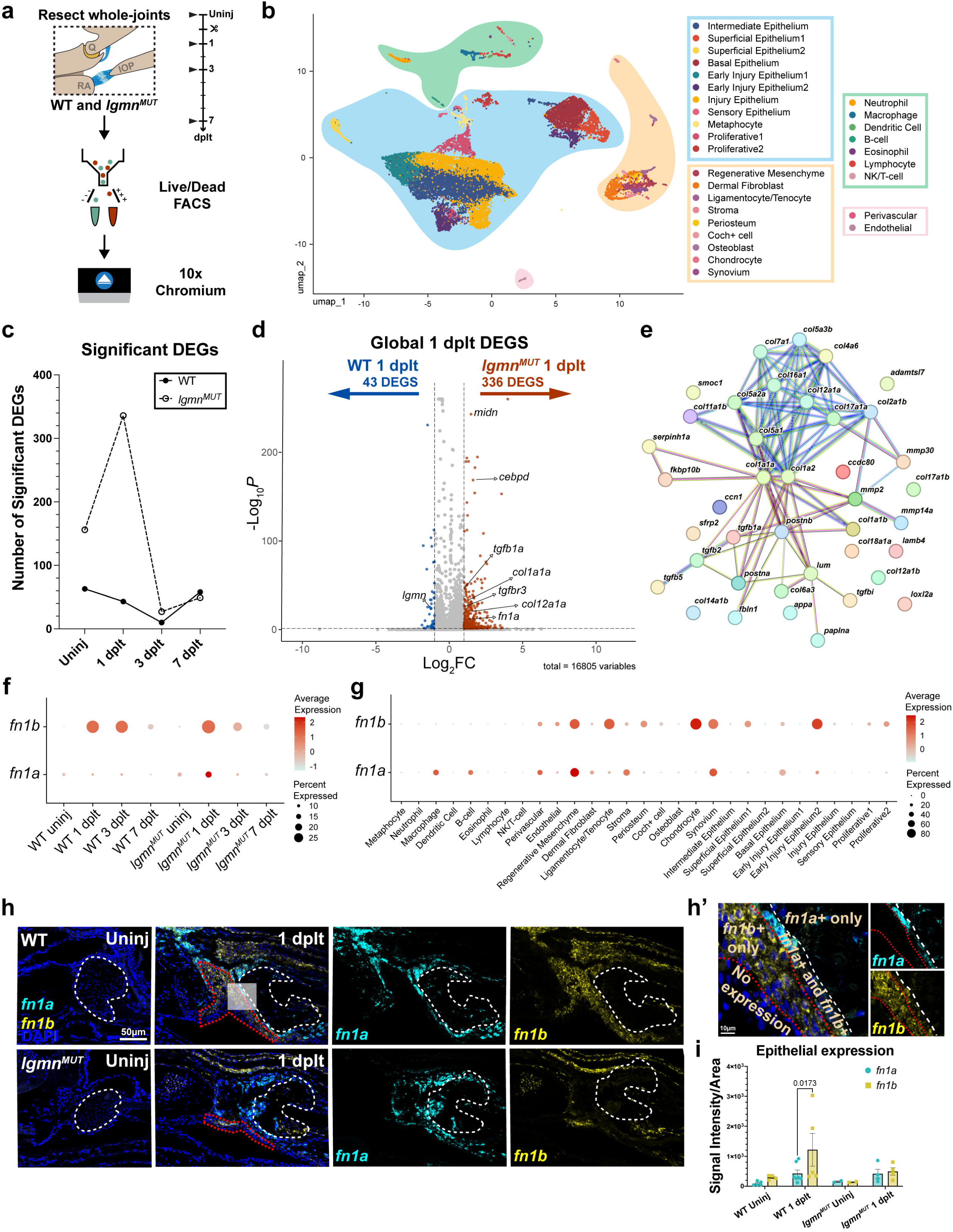
Transcriptional response after ligament injury. a) Overview workflow for scRNAseq data collection and processing. b) Visualization of clusters in WT and *lgmn^MUT^* at all timepoints in UMAP. Clusters were broadly categorized by cell type: immune (green), endothelium (pink), epithelium (blue), and skeletal (peach). c) Number of significant DEGs identified for each genotype when comparing global transcriptional response of WT vs. *lgmn^MUT^* of the same timepoint. d) Volcano plot of significant DEGs identified globally at 1 dplt between WT and *lgmn^MUT^*. e) STRING diagram of the ECM organization protein-network identified enriched in *lgmn^MUT^* at 1 dplt. (f, g) Average and percent expression profile of *fn1a* and *fn1b* by timepoint and genotype (f), and by cell type (g). (h) RNAscope of *fn1a* (cyan), *fn1b* (yellow), and counterstained with DAPI (blue) in WT and *lgmn^MUT^* uninjured and 1 dplt. (h’) Inset of transparent white box in (h) map of expression pattern. White dashed-line marks ligament. Red dotted line is epithelial ROI quantified in (i). (n=3-5 per genotype) (i) Quantification of *fn1a* and *fn1b* RNAscope signal in epithelial region denoted in (h). Error bars = SD. Scale bars = 50µm (h), 10um (h’).

To identify differences in global transcriptional responses, we compared WT and *lgmn^MUT^* to determine significant differentially expressed genes (DEGs) at each timepoint (pvalue <= 0.05;LFC±1) (Table S2). The timepoint with the largest transcriptional difference was 1 dplt with 43 enriched DEGs in WT and 336 enriched DEGs in *lgmn^MUT^* (Figure 2c, d). *lgmn* was among the upregulated DEGs in WT, suggesting that absence of Lgmn protein in *lgmn^MUT^* may influence *lgmn* mRNA expression. Analysis of STRING-db networks of WT and *lgmn^MUT^* DEGs showed a minor enrichment for methylarsonate reductase and glutathione dehydrogenase activity in WT at 1 dplt (Figure S1c, Table S3). Interestingly, the enriched pathways identified in *lgmn^MUT^* included ECM-related processes, cell adhesion, peptidase regulator activity, and type II TGFβ receptor binding (Figure S1d, Table S3). Consistent with a dysregulated ECM remodeling response observed in *lgmn^MUT^*, there was upregulation of transcription of collagens including *col1a1a, col5a3b,* and *col12a1a*, as well as Tgfβ signaling factors, including *tgfb1a, tgfb2, tgfb5, tgfbi,* and *tgfbr3* (Figure S2e). Taken together, this suggests altered Tgfβ activity may be underlying excessive ECM expression in *lgmn^MUT^*, which is characteristic of a pathological fibrotic response.

### Transcriptional regulation of *fn1a* and *fn1b* expression are altered in *lgmn^MUT^*

Increased fibronectin 1 (FN1) is commonly associated with a fibrotic response, and *lgmn* has been shown to alter *Fn1* expression through Tgfβ signaling^35,36^. Zebrafish have two copies of fibronectin due to a genome duplication: *fn1a* and *fn1b,* and there is increasing evidence for diverging functions for these paralogs^37^. We sought to examine the differential roles of *fn1a* and *fn1b* in the context of tissue regeneration. scRNAseq analysis showed *fn1a* and *fn1b* expression peaked at 1 and 3 dplt in both WT and *lgmn^MUT^* (Figure 2f). High expression of *fn1a* and *fn1b* were in mutually exclusive cell types, except within the regenerative mesenchyme and synovium clusters, which had equivalent expression of both transcripts (Figure 2g). This cell type-dependent expression suggests non-redundant function and differential regulation of expression determined by tissue type.

To identify if this cell type-specific expression pattern was spatially distinct *in vivo,* RNAscope was used to visualize *fn1a* and *fn1b* expression domains. In agreement with our scRNAseq expression profile, we found *fn1a* and *fn1b* have spatially discrete expression domains. Uninjured ligaments had low *fn1b* expression and little to no *fn1a* expression. By 1 dplt, there was increased expression of both *fn1a* and *fn1b* in WT and *lgmn^MUT^* (Figure 2h). The expression pattern showed a distinct spatial arrangement of cell layers that were either *fn1a+* or *fn1b+* with a minimal overlap of a monolayer of *fn1a+;fn1b+* double positive cells sandwiched between the expression domains (Figure 2h’). WT and *lgmn^MUT^* displayed the same organization at 1 dplt, but *lgmn^MUT^* have less epithelial *fn1b* expression (Figure 2i). These data suggest the cytoarchitecture of *fn1a* and *fn1b* expression are used to preserve tissue organization during regeneration.

### Injury drives a shift in proteomic landscape and factors involved in ECM remodeling are dysregulated in *lgmn^MUT^*

Current research in regenerative biology is dominated by epigenetic and transcriptomic assays that provide a genetic blueprint about mRNA-level changes but ultimately is not a sufficient proxy for complete understanding of biological function at the protein level. To address this gap, we used mass-spectrometry based proteomics to identify how the functional molecular landscape changes to support cellular processes that lead to either a regenerative or scarring outcome. We used label-free liquid chromatography-mass spectrometry (LC/MS) to profile the proteomic signature of uninjured contralateral and injured 3 dplt joints from WT and *lgmn^MUT^* (Figure 3a). Data-independent acquisition was used as an unbiased method to detect a broad spectra of peptides. After QC filtering, we identified 9,042 unique proteins across the four experimental conditions and PCA analysis using the top 300 variable proteins showed that PC1 (31.3%) separated by injury state and PC2 (19.4%) separated by genotype (Figure 3b). These data demonstrate that injury state has a larger influence in proteome composition than genotype.

**Figure 3.**
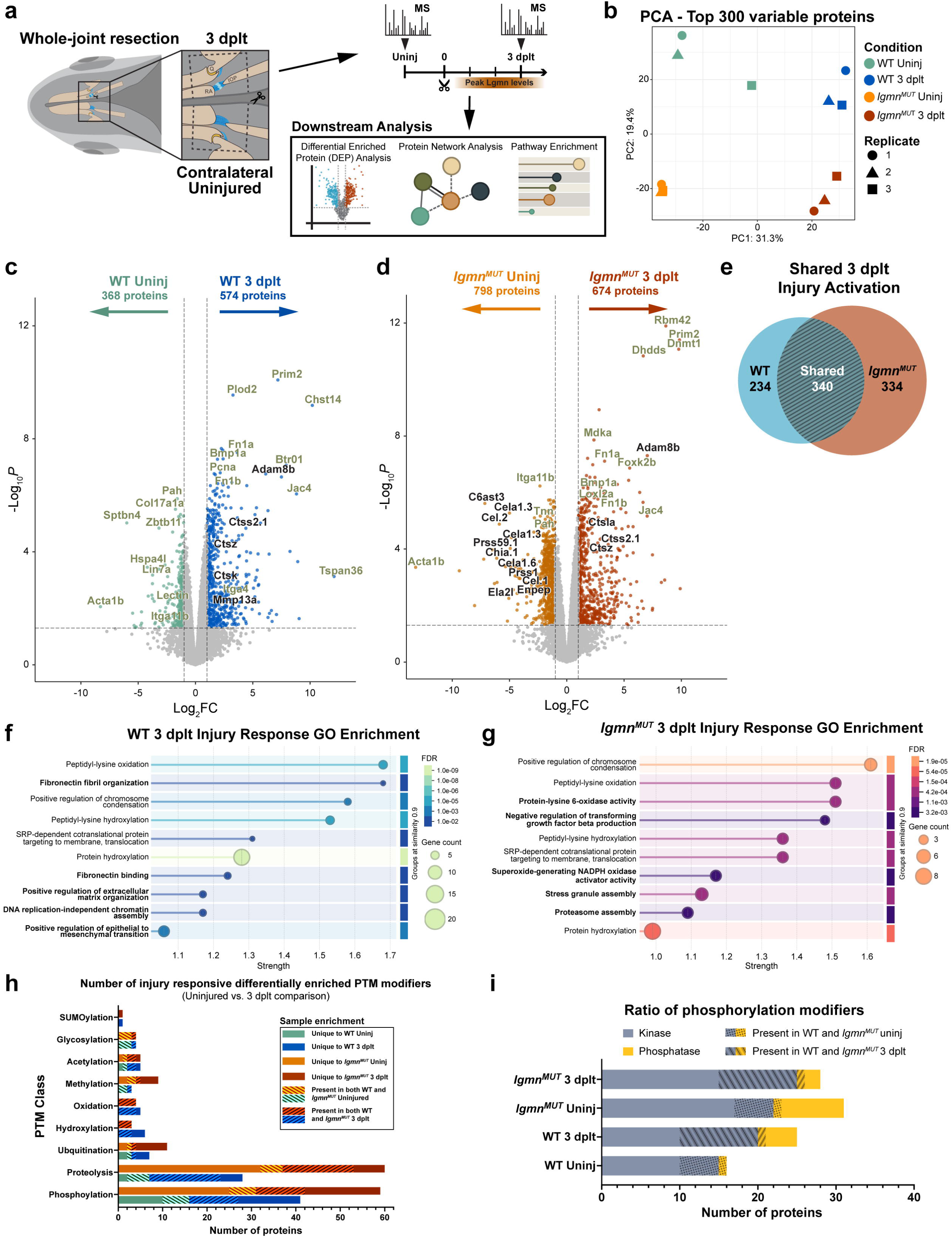
Mass spectrometry proteomic profiling of regenerate tissue reveal enrichment of post-translational modifiers. a) Workflow of sample collection for LC/MS proteomics and downstream analysis. b) Principal component analysis of the top 300 variable proteins. c) Volcano plot of significant DEPs in WT injury response identified by uninjured vs. 3 dplt comparison. Proteins in black represent proteases. d) Volcano plot of significant DEPs identified in *lgmn^MUT^* injury response identified by uninjured vs. 3 dplt comparison. Proteins in black represent proteases. e) Venn-diagram representing the overlap of shared injury-activated proteins identified by comparing list of 3 dplt enriched for either WT or *lgmn^MUT^* from (c) and (d). f) Enriched GO terms of WT injury-activated proteins detected in (c). Bolded terms are specific to WT injury-activated proteins. g) Enriched GO terms of *lgmn^MUT^* injury-activated proteins detected in (d). Bolded terms are specific to *lgmn^MUT^* injury-activated proteins. h) Number of injury responsive DEP shown in (c) and (d) classified as post-translational modifiers. No patterned bars indicate proteins unique to that sample. Left-leaning shaded bar indicates the number of proteins enriched in both WT and *lgmnMUT* uninjured and right-leaning shaded bar indicates enrichment in both genotypes at 3 dplt. i) Ratio of kinase and phosphatase phosphorylation factors in (h) split by sample.

To determine proteomic response to injury and how it differs between genotypes, we ran differentially enriched protein (DEP) analysis to compare uninjured verses 3 dplt samples for WT or *lgmn^MUT^* . We found that 574 proteins are significantly enriched in WT 3 dplt compared to 368 enriched in uninjured WT (Figure 3c, Table S4). In *lgmn^MUT^,* we observed 798 enriched proteins in uninjured states and 674 significantly enriched in *lgmn^MUT^* 3 dplt (Figure 3d, Table S5). These lists comprised the injury-activated proteins in WT and *lgmn^MUT^* and 340 proteins were shared between both genotypes. There were also proteins uniquely enriched in each genotype, 234 are WT injury-specific and 334 are *lgmn^MUT^* injury-specific (Figure 3e, Table S6). Shared biological processes in WT and *lgmn^MUT^* were determined by performing protein network analysis using STRING and Gene Ontology (GO). We observed enrichment for PTM regulation processes such as peptidyl-lysine modification, peptidyl-proline modification, and peptidyl-amino acid modification in both genotypes (Figure 3f, g, Table S7, S8). While there was a similar broad injury response shared by both genotypes, we also identified differences between the WT and *lgmn^MUT^*response. Interestingly, processes such as fibronectin binding and fibril organization, positive regulation of ECM organization, DNA replication-independent chromatin assembly, and positive regulation of epithelial to mesenchymal transition were specifically enriched in WT samples at 3 dplt and not in *lgmn^MUT^* (Figure 3f, bolded, Table S7). Processes that were specific to *lgmn^MUT^* only included negative regulation of Tgfβ production, superoxide-generating NADPH oxidase activator activity, stress granule assembly, and proteasome assembly (Figure 3g, bolded, Table S8). These results show a shared injury response of upregulated ECM crosslinking through peptidyl-lysine oxidation and hydroxylation. However, processes involving fibronectin binding and organization were enriched in WT but not *lgmn^MUT^*, consistent with Fn1a/b as a known Lgmn substrate^38,39^.

Manual identification of common post-translational modifiers among the DEP identified that proteins involved in phosphorylation and proteolysis were the most abundant post-translational modifiers upregulated after injury (Figure 3h). Proteins involved in glycosylation were present in both the WT and *lgmn^MUT^* 3 dplt, indicating that glycosylation is similar in both regenerates. Histone-modifying methyltransferases (Prmt1, Prmt6, Ntmt1) were enriched only in *lgmn^MUT^*3 dplt samples, suggesting dysregulation in epigenetic remodeling may occur in *lgmn^MUT^*(Figure 3h). Separating phosphorylation factors into kinases or phosphatases showed *lgmn^MUT^*uninjured samples had more enrichment for phosphatases compared to WT uninjured (Figure 3i). Cyclin-dependent kinases (Cdk1, Cdk2, Cdk13) were enriched only in *lgmn^MUT^*3 dplt samples, which may indicate dysregulation of cell-cycle that contributes to the decreased rates of proliferation previously observed^7^. These data indicate dynamic regulation of post-translational modifiers in the regenerative proteome.

### Immune and skeletal cell populations drive post-translational modifications

We leveraged both transcriptomic and proteomic modalities to deconvolve proteomic data into cell type-associated functional modules. This analysis associates expressed proteins with transcriptional expression at single cell resolution. Pairing these modalities bridges the gap between transcriptional response and functional output. Here, we couple these modalities to identify molecular signatures, infer functionality, and trace cellular origin. To link the bulk-proteomics with scRNAseq, a list of 8,211 genes found in both datasets were used to guide non-negative matrix factorization (NMF) (Figure 4a). NMF was applied to each sample with multiple parameter values (K=5-15) that identified 15 consensus module programs (MP). After QC filtering, we identified 10 MPs ranging from 2 to 60 genes. Cells within each module grouped along cell identity that are broadly classified as epithelial, immune, skeletal, and proliferative cell identities (Figure 4b).

**Figure 4.**
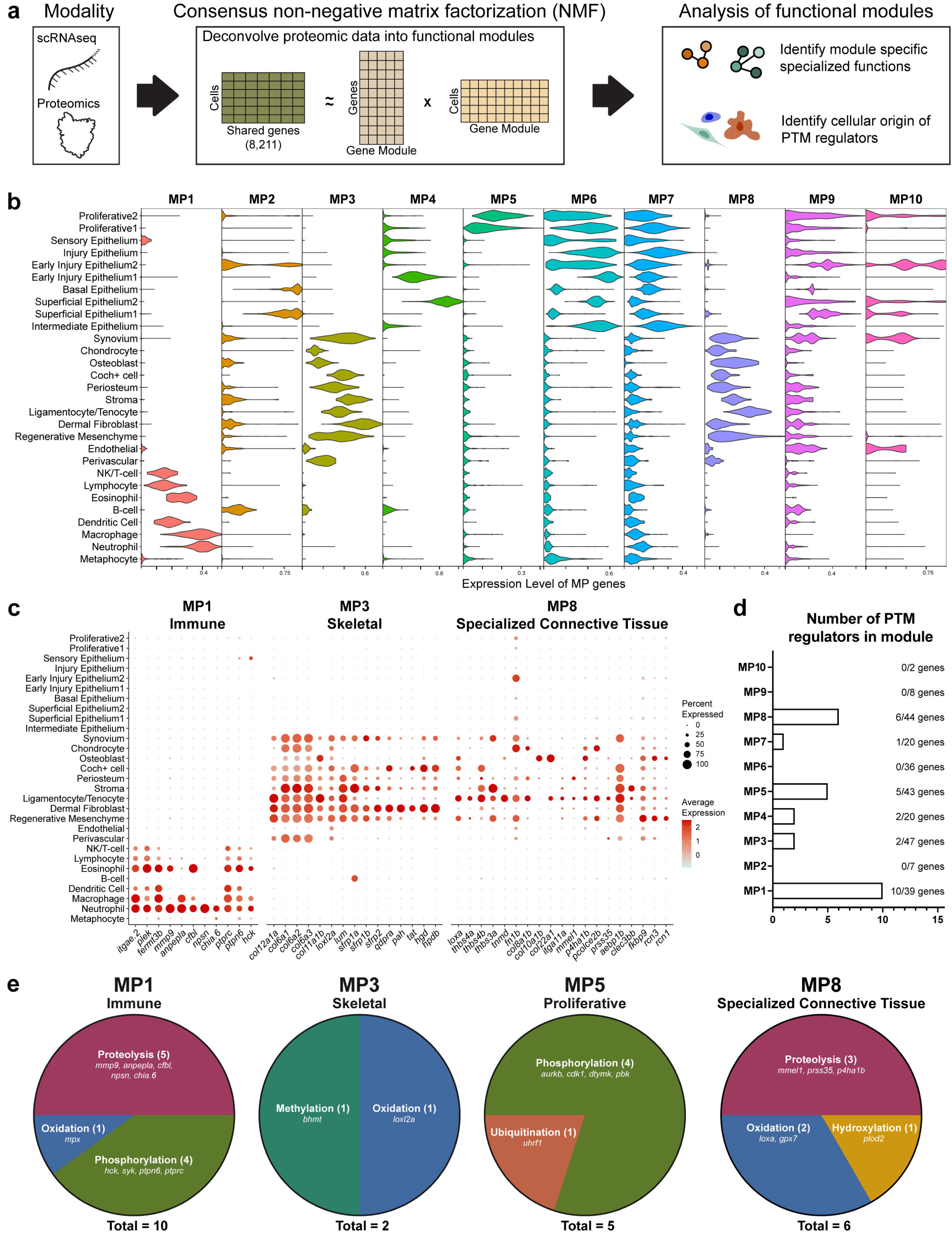
Multimodal NMF show immune and skeletal populations are main cellular drivers of post-translational modifications. a) Workflow of multimodal NMF analysis using bulk proteomics and scRNA sequencing. b) Violin plot of highest scoring cells for each gene module program. c) Dot plot of key driving genes for MP1, MP3, and MP8 split by cell type. d) Number of post-translational modifying proteins driving each module. e) Ratio of PTM classes identified in each module.

Immune cells were enriched in MP1 and GO analysis of the driving genes for MP1 included integrin-signaling (*itgae.2, plek, fermt3b)* and proteolytic proteins (*anpepla, cfbl, mmp9, npsn, chia.6*) (Figure 4c). Neutrophils and macrophages were highest expressors of these proteases, indicating that they are a primary source of proteases during early ligament regeneration. Skeletal populations clustered into either MP3 or MP8, of which MP8 represented highly specialized cells specific to ligament tissue. GO analysis of MP3 genes included ECM components (*col12a1a, col6a1, col6a2, col6a3, col11a1b, loxl2a, lum),* modulators of Wnt signaling (*sfrp1a, sfrp2a, sfrp1b),* and tyrosine metabolism (*qdpra, pah, tat, hgd, hpdb*) (Figure 4c). A subpopulation of “specialized connective tissue” skeletal cells including regenerative mesenchyme, ligamentocytes/tenocytes, osteoblasts, and chondrocytes scored highest for MP8. Characterized by expression of tissue-specific structural ECM genes (*loxa, thbs4b, thbs4a, tnmd, fn1b,* and multiple collagens), matrix remodeling proteases (*mmel1, p4ha1b, prss35)*, cell-ECM interaction (*itga11a),* and calcium ion binding (*clec3bb, fkbp9, rcn1, rcn3)* (Figure 4c). MP8 represents a functional class of specialized cells maintaining connective tissue structure through ECM remodeling and regulation of calcium signaling in response to external stimuli.

Using the driving genes for each module, we next identified the class of post-translational modifications that were employed within each broad cell identity class. PTM regulators involved in proteolysis, phosphorylation, oxidation, methylation, acetylation, and ubiquitination were identified (Figure 4d). Immune (MP1) and specialized connective tissue (MP8) modules had the highest percentage of PTM regulators with 25.6% (10/39 genes) and 13.6% (6/44 genes), respectively (Figure 4d). Proteolytic and phosphorylation proteins were among the most common class of PTMs (Figure 4e). A clear link between post-translational modifiers and their corresponding function can be seen within MP5 which represents proliferative cells. Here, we find cell cycle progression is regulated through phosphorylation with *aurkb*, *cdk1*, *pbk,* and *dtymk* (Figure 4e). Proteases were a large class of post-translational modifiers in MP1 (*mmp9, anpepla, cfbl, npsn, chia.6*) and MP8 (*mmel1, prss35, p4ha1b*), showing that immune and a specific subset of skeletal cells are the primary cellular contributors of proteases into the regenerate.

### Lgmn deficiency impairs proper Fn1a/b network organization

We next sought to identify how the ECM was remodeled in response to the influx of proteolytic proteins into the regenerate. Fn1a/b is a known substrate of Lgmn and it is influential in establishing tissue organization through guiding future collagen deposition^40,41^. We investigated how Lgmn deficiency impacts Fn1a/b abundance and organization during regeneration. To determine how Fn1a/b abundance changed *in vivo*, we used western blot analysis of whole-joint tissue lysates collected from WT and *lgmn^MUT^* at uninjured, 1, 3, and 7 dplt. Since the anti-Fn1 antibody detects both Fn1a and Fn1b, we could not identify which bands belonged to either the a or b isoform. To measure intact Fn1a/b, 270kDa+ bands were considered fully mature fibronectin^42^ (Figure 5a). We observed an injury-specific upper band at the 1 and 3 dplt timepoints that waned by 7 dplt (Figure 5a). Comparable levels of 270kDa+ Fn1a/b were detected in WT and *lgmn^MUT^*throughout the time course of healing (Figure 5b). We next assessed Fn1a/b fibril network organization during healing. IF staining was used to visualize the spatial organization of Fn1a/b throughout the regenerate tissue in WT and *lgmn^MUT^* at uninjured, 1, 3, 7, and 14 dplt (Figure 5c). These data showed dynamic remodeling of the Fn1a/b network throughout the course of healing in both WT and *lgmn^MUT^.* By 14 dplt, the Fn1a/b network in WT begins to resemble the densely packed network seen in uninjured ligaments, while it is disorganized and has a patchy pattern in *lgmn^MUT^*.

**Figure 5.**
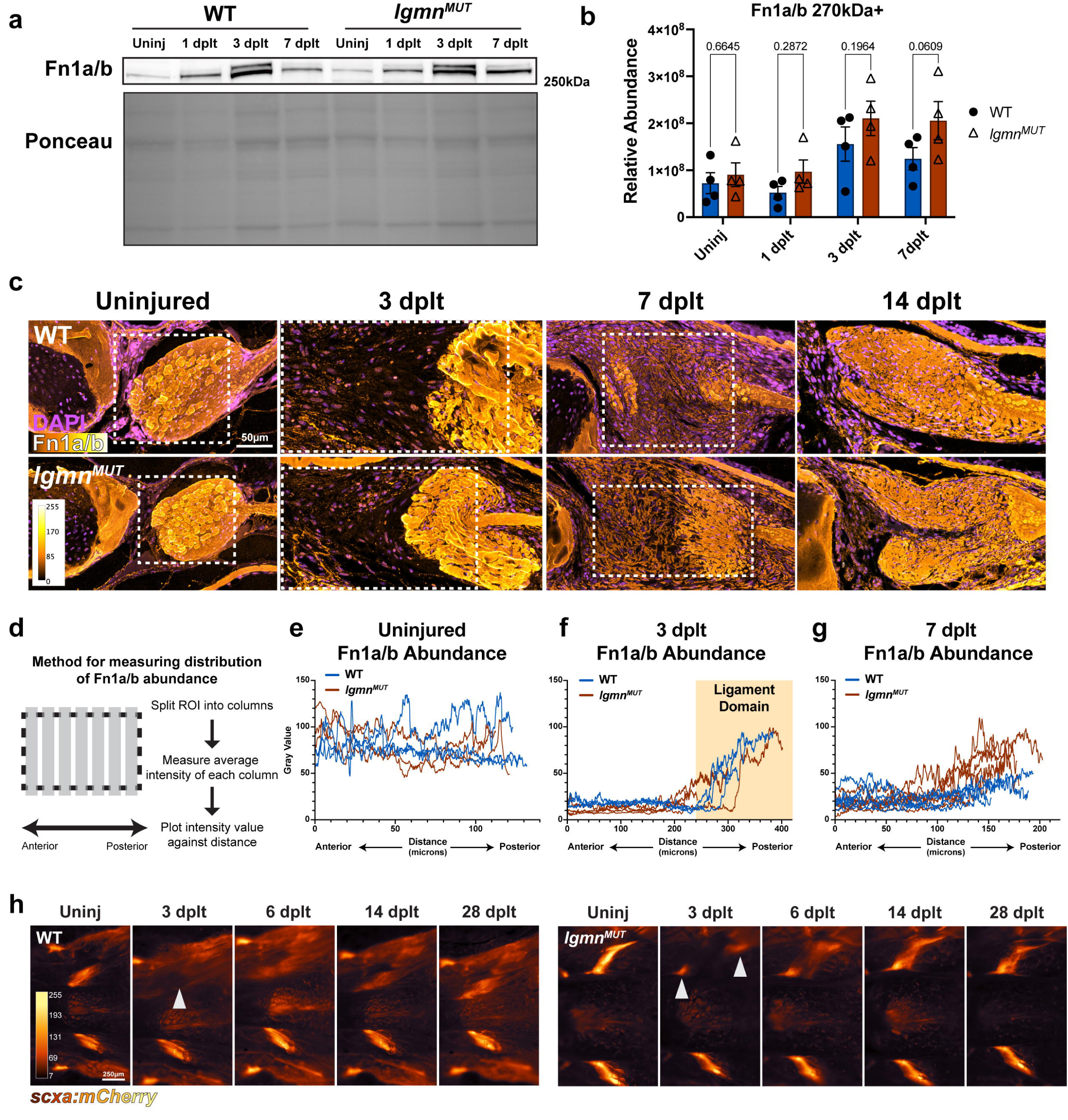
Fibronectin 1a/b network remodeling is impaired in *lgmn^MUT^*. (a) Fn1a/b western blot of whole joints collected from WT and *lgmn^MUT^* uninjured, 1, 3, and 7 dplt (n=4 per timepoint). High molecular weight bands 270kDa+ represented fully mature Fn1a/b. (b) Quantification of Fn1a/b 270kDa+ bands detected in (a) western blot relative to total protein input. (n=4-5 per timepoint). Error bars = SD. (c) Immunofluorescence of Fn1a/b in uninjured, 3, 7 and 14 dplt in WT and *lgmn^MUT^* (n=5-10 per timepoint). Nuclei (magenta) and Fn1a/b (orange hot) are pseudo colored to represent increased signal intensity. White dashed-square box is quantification ROI in (g-i). (d) Schematic of method to measure distribution of Fn1a/b abundance along tissue distance from anterior to posterior. (e-g) Plots of Fn1a/b abundance along an anterior to posterior axis of ROI in (c) from uninjured (n=3 per genotype) (e), 3 dplt (n=3) (f), and 7 dplt (n=5-6) (g). Ligament domain at 3 dplt was determined by measuring the distance from edge of ligament to IOP bone. (j) Repeated live imaging of *scxa:mCherry* expression in WT and *lgmn^MUT^* at uninjured, 3, 6, 14, and 28 dplt (n=3-4 per genotype). Arrowheads indicate ligament tissue. Scale bar = 50µm (e) and 250µm (h).

Failure to break down the collagen within the injured ligament bundle was observed in *lgmn^MUT^* at 3 dplt^7^. To show a requirement for Lgmn in spatially regulating this tissue remodeling, we quantified the spatial abundance of Fn1a/b relative to distance throughout the tissue. The average signal intensity of Fn1a/b was measured along an anatomical Anterior/Posterior axis by drawing a region of interest (ROI) from the retroarticular process (RA, anterior) to the tip of the interopercular bone (IOP, posterior), dividing into columns, and using the plot profile function in ImageJ to measure the average intensity of each column (Figure 5d). Uninjured ligaments have uniform and densely packed Fn1a/b (Figure 5e). The preexisting dense Fn1a/b network was easily identified at 3 dplt where the injured ligament sits adjacent to the newly forming Fn1a/b network of the granulation tissue (Figure 5f). Quantification at 3 dplt showed the intensity spike at the posterior end was due to the injured ligament, noted as the ligament domain (Figure 5f). These results showed a stark change in tissue topography at 3 dplt, where areas of low Fn1a/b sat adjacent to areas of high Fn1a/b (Figure 5f). By 7 dplt, Fn1a/b abundance was uniform in WT, but *lgmn^MUT^* retained the trend of increased abundance near the posterior end and had a magnitude of intensity comparable to 3 dplt (Figure 5g). Together, these data suggest that *lgmn^MUT^* fail to break down the Fn1a/b network and reabsorb the ligament from 3 to 7 dplt, with this initial mis-patterning of Fn1a/b likely influencing downstream tissue organization and cellular response to the altered microenvironment.

### Ligamentocyte dedifferentiation is impaired in *lgmn^MUT^*

Unlike mammalian ligament healing, post-mitotic mature ligamentocytes in adult zebrafish undergo dedifferentiation following ligament transection^7^. To determine how the ligamentocytes responded to the impaired ECM remodeling in the ligament bundle, we visualized ligamentocyte fate by repeated live imaging of *scxa:mCherry* after ligament transection. At 3 dplt, *scxa:mCherry* is completely absent in the injured area in WT (n=4) but persistent in the ligament bundles in *lgmn^MUT^*(n=4/6) (Figure 5h). This failure to turn off expression of the mature ligamentocyte marker *scxa* suggests that proper ECM remodeling is required for ligamentocyte dedifferentiation.

### Protease diversity is decreased after injury in *lgmn^MUT^*

To determine how the regenerate landscape differed between WT and *lgmn^MUT^,* the proteomic composition of WT 3 dplt and *lgmn^MUT^*3 dplt were compared to identify genotype-driven DEPs. This comparison identified 113 proteins significantly enriched in WT 3 dplt and 117 proteins in *lgmn^MUT^* 3 dplt (Figure 6a, Table S9. GO analysis of STRING-db networks showed WT 3 dplt samples enriched for processes involving peptidase activity, proteolysis, cell-matrix interactions, and immune response (Figure 6b, Table S10). Protein networks of *lgmn^MUT^* 3 dplt samples were enriched for processes including gas transport, oxygen binding, heme binding, and hydrogen peroxide catabolic process (Figure 6c, Table S10). In comparison to WT 3 dplt regenerates, *lgmn^MUT^* lost enrichment for pathways associated with a wound response such as immune response, cell-matrix adhesion, and proteolysis. Many of the WT significantly expressed proteins at 3 dplt were identified within the proteolysis network (Figure 6d). Interestingly, in addition to complete loss of Lgmn in both *lgmn^MUT^* conditions, the proteases Cela 1.3, C6ast3, and Prss59.1 were not detected by LC/MS in all three *lgmn^MUT^*3 dplt replicates (Figure 6e). These proteases are present in uninjured *lgmn^MUT^* at similar levels to WT uninjured and 3 dplt but plummet in response to injury in *lgmn^MUT^*. Indicating that their absence in *lgmn^MUT^* at 3 dplt is not due to an inability to synthesize these proteases, rather it is due to an injury-specific response. To determine global levels of proteases present in the LC/MS, we aggregated the intensity readings of all proteases to measure total abundance. Out of the 9,042 proteins, we identified 265 proteases across all samples and global protease abundance was quantified by aggregating their intensity readings. Global protease abundance was significantly decreased in *lgmn^MUT^* compared to WT at 3 dplt (p=0.0366) (Figure 6f) and this decrease was not due to a decrease in total protein (p=0.6869) (Figure 6g). These data show a requirement for Lgmn in regulating global protease levels after injury and suggest that decreased protease levels in *lgmn^MUT^* may impair proteolytic activity.

**Figure 6.**
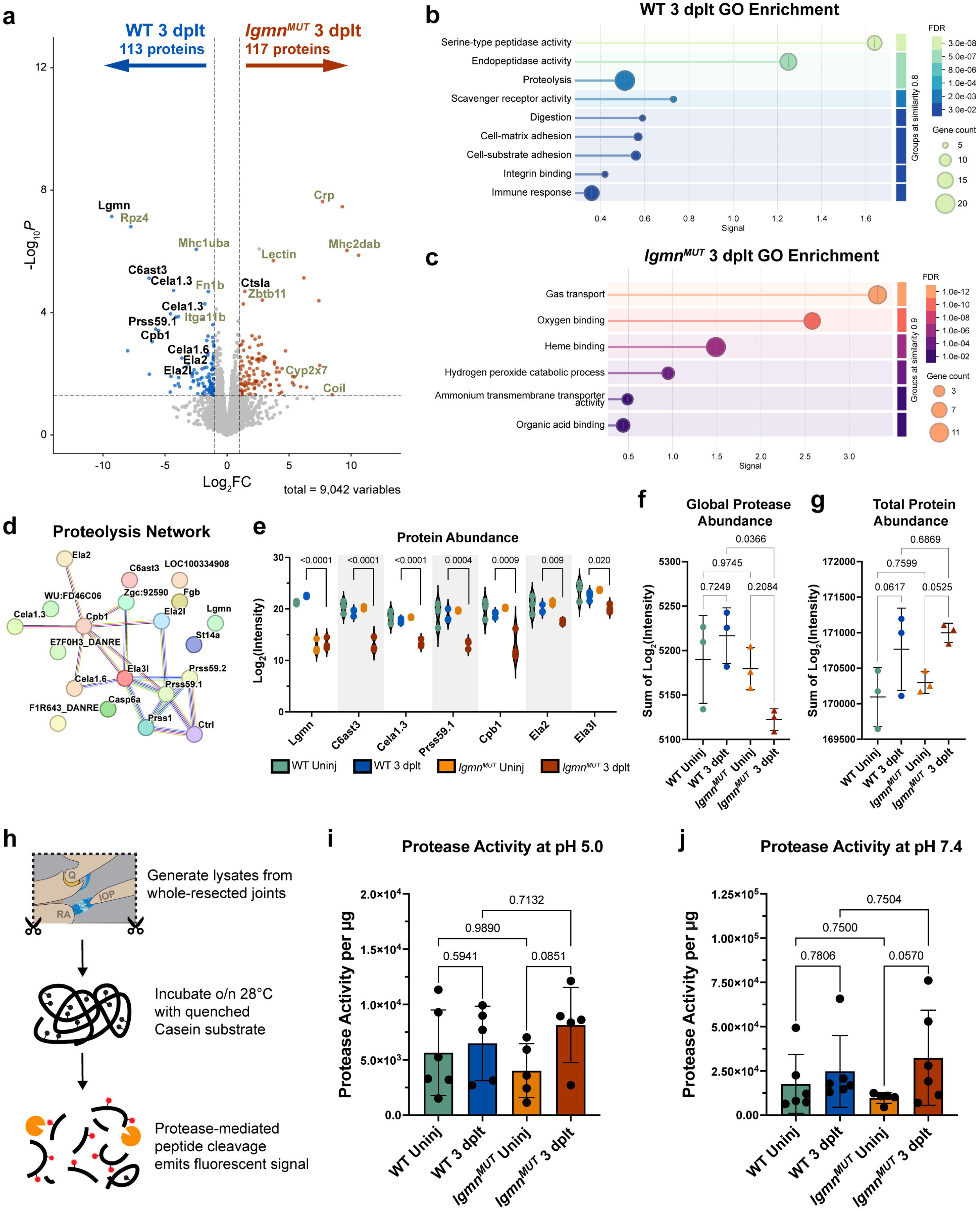
Global protease abundance decreases after injury in *lgmn^MUT^* but proteolytic activity is unaltered. a) Volcano plot of significant DEPs between WT 3 dplt (left, blue) and *lgmn^MUT^* 3 dplt (right, red). Proteases are marked in black text. b) Enriched GO terms of upregulated proteins in WT at 3 dplt. c) Enriched GO terms of upregulated proteins in *lgmn^MUT^* at 3 dplt. d) Proteolysis STRING network enriched in WT at 3 dplt. Cyan and magenta lines represent known interactions. Green, red, and blue lines represent predicted interactions. Chartreuse, black and periwinkle lines represent text mining, co-expression, and protein homology interactions. e) LC/MS abundance of proteases in proteolysis network. f) Global protease abundance determined by total sum of log-transformed intensity readings of all proteases present in each sample detected by LC/MS. Error bars = SD. g) Schematic of workflow used to measure global protease activity. h) Quantification of protease activity measured normalized to protein input under acidic conditions at pH 5.0 (n=5-6 per timepoint). Error bars = SD. i) Quantification of protease activity measured normalized to protein input under neutral conditions at pH 7.4 (n=5-6 per timepoint). Error bars = SD.

### Global protease activity is unaltered in *lgmn^MUT^*

To determine if decreased protease abundance in *lgmn^MUT^*had a functional consequence, we measured global protease activity using the EnzChek Protease Assay, which detects activity from a broad range of proteases. Protease-catalyzed hydrolysis of a casein substrate emits a fluorescent signal used as a quantitative measurement for protease activity. Fresh whole-joint lysates were generated from uninjured and 3 dplt WT and *lgmn^MUT^* and incubated with the casein substrate at either an acidic pH of 5.0, optimal for lysosomal proteases, or neutral pH of 7.4, optimal for most proteases (Figure 6h). Contrary to our expectations, these results showed similar protease activity levels in WT and *lgmn^MUT^* at baseline and after injury under both acidic and neutral conditions (Figure 6i, j). Furthermore, we observed a modest increase of protease activity in WT after injury under neutral (p=0.7806) and acidic (p=0.5941) conditions. Unexpectedly, *lgmn^MUT^* had a more significant increase in global protease activity after injury under acidic (p=0.0851) and neutral (p=0.0570) conditions (Figure 6H, I). These results were unexpected, as we found similar levels of global protease activity between WT and *lgmn^MUT^*at 3 dplt. Moreover, *lgmn^MUT^* experienced a large spike in proteolytic activity in response to injury in comparison to the more modest increase observed in WT when comparing changes from uninjured to 3 dplt.

### Lgmn is required for the concentrated localization of Ctsla at the injury edge

The increase of protease activity in *lgmn^MUT^* at 3 dplt was surprising given the reduced protease abundance detected by LC/MS. We hypothesized that this was due to a compensatory response to Lgmn deficiency in *lgmn^MUT^*. To determine which proteases were compensating for the loss of Lgmn, we used LC/MS to identify proteases enriched in *lgmn^MUT^* at 3dplt. Ctsla, Ctsh, Napsin-A (Napsa), Granzyme-like protein 1, and Duodenase-1 were significantly increased in *lgmn^MUT^* compared to WT at 3 dplt (p-value <= 0.05, Log_2_FC >=0.5) (Table S9). Interestingly, three of the five upregulated proteases are classified as lysosomal proteases (Ctsla, Napsa, and Ctsh). Among other cathepsins identified, only Ctsla was significantly increased in *lgmn^MUT^*compared to WT at uninjured baseline levels (p=0.0418) and 3 dplt (p=0.0026) (Figure 7a). At the transcript level, macrophages had the highest expression of *ctsla* and *ctsh,* but *ctsla* was also expressed by a broader range of cell types (Figure 7b). Ctsla localization was visualized with IF, which confirmed increased expression of Ctsla diffusely throughout the uninjured ligament in *lgmn^MUT^* compared to minimal expression in WT (Figure 7c). Interestingly, high concentrations of Ctsla were observed at the injury interface at 3 dplt in WT but not in *lgmn^MUT^* (Figure 7c’). Indeed, quantification of Ctsla at the integration site was increased after injury in WT but not in *lgmn^MUT^* (Figure 7c’ (red dotted line), 7d). We found both Lgmn and Ctsla localized around the ligament injury border and previous work has shown Lgmn drives Ctsl processing in murine cells^7^. To determine where Ctsla and Lgmn were spatially interacting, Ctsla was visualized with tsa multiplexed with Lgmn IF. High-resolution confocal microscopy showed that Lgmn and Ctsla were co-localized throughout the injury and were highly concentrated in cells surrounding the injured ligament in WT (Figure 7e, e’). RNAscope showed that *ctsla* and *lgmn* were co-expressed in the same cells, specifically in *mpeg1.1+* macrophages (Figure 7f). Taken together, these data suggest that high concentrations of Ctsla and Lgmn near the injury edge are produced by macrophages, and their subcellular co-localization suggests that Lgmn may be involved in Ctsla processing.

**Figure 7.**
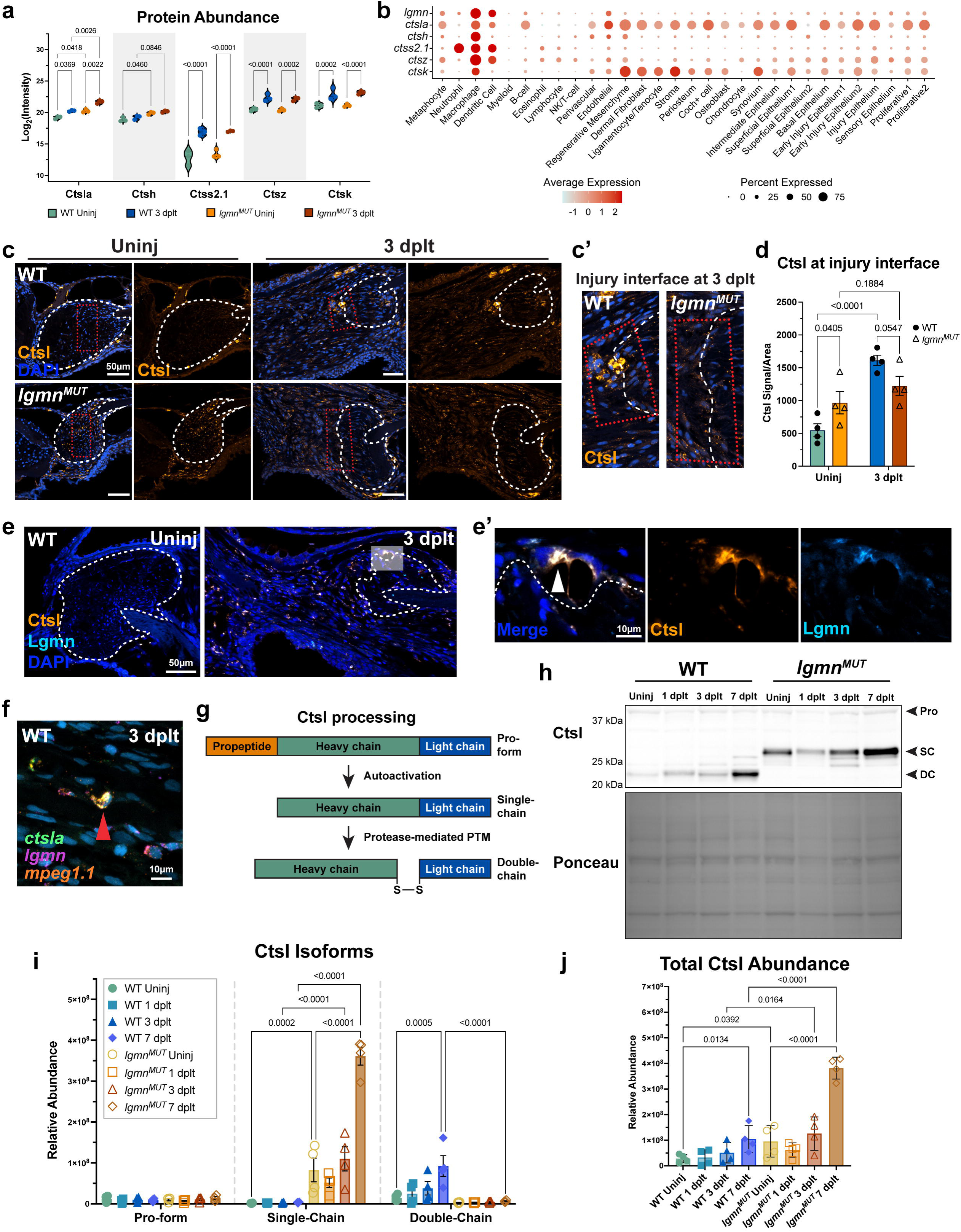
Increased Cathepsin La abundance and altered processing in *lgmn^MUT^*. a) Protein abundance of cathepsins significantly upregulated in response to injury detected by LC/MS. b) Expression profile of *lgmn* and cathepsins by cell type in scRNAseq. c) IF for Ctsla (orange) in WT and *lgmn^MUT^* uninjured and 3 dplt. White dashed-line marks ligament. Red dotted line is ROI drawn around the injury interface quantified. (c’) Inset of injury interface quantified in (d). d) Quantification of Ctsla abundance at the injury interface in (c) (n=4 per timepoint). e) Max projection of Lgmn (cyan) and Ctsla (orange) co-localization *in vivo* visualized with Ctsla-tsa and Lgmn-IF counterstained with DAPI (blue) in WT at uninjured and 3 dplt. White dashed-line marks ligament. (e’) Inset of area marked by a white transparent box in (e), showing a single z-slice of high concentration of Ctsla and Lgmn co-localized at injury interface. f) RNAscope of *mpeg1.1*+ (orange) macrophage co-expressing *lgmn* (magenta) and *ctsla* (green), nuclei counterstained with DAPI in teal. g) Diagram representing Ctsl proteolytic processing to the catalytically active single and double-chain forms. h) Western blot of Ctsl in WT and *lgmn^MUT^* uninjured, 1, 3, and 7 dplt lysates generated from whole-resected joints (n=4 per timepoint). Pro = pro-form, SC = Single-chain, DC = double-chain. i) Quantification Ctsl abundance separated by pro-form, SC, and DC isoforms detected by WB in (h) relative to total protein input. Error bars = SD. j) Quantification of total Ctsl abundance by WB in (h) relative to total protein input. Error bars = SD. Scale bars = 50µm (c, e), 10µm (e’, f)

### Lgmn drives Ctsla processing into its double-chain form during regeneration

Ctsla is processed from its inactive pro-form to its active single-chain (SC-Ctsla) form through autoactivation and is further proteolytically processed into its active double-chain (DC-Ctsla) form (Figure 7g). Here, we examined Ctsla processing in zebrafish and assessed *lgmn^MUT^* for alterations in processing. To confirm that *lgmn^MUT^* have increased abundance of Ctsla *in vivo*, total Ctsla abundance was determined through western blot analysis from whole-joint lysates collected from WT and *lgmn^MUT^*at uninjured, 1, 3, and 7 dplt timepoints (Figure 7h). DC-Ctsla was dominant in WT and is significantly elevated at 7 dplt compared to uninjured levels (Figure 7i). Interestingly, SC-Ctsla is the dominant species present in *lgmn^MUT^*, and no DC-Ctsla was detected (Figure 7i). SC-Ctsla was significantly elevated in *lgmn^MUT^*compared to WT at uninjured, 3, and 7 dplt (p=0.0002, <0.0001, <0.0001, respectively) (Figure 7i). Furthermore, WB confirmed *lgmn^MUT^*have increased levels of Ctsla at a baseline uninjured state (p=0.0134) as well as after injury at 3 and 7 dplt compared to WT (p=0.0164 and <0.0001, respectively) (Figure 7j). These data suggest a potential compensatory upregulation of Ctsla in *lgmn^MUT^* and demonstrate that Lgmn is required for processing SC-Ctsla into DC-Ctsla during homeostasis and tissue injury response. Taken together, this work demonstrates an important role for protease-mediated post-translational modifications for scarless ligament regeneration.

## Discussion

Fibrotic scarring often impairs organ function and is common after tissue injury in adult mammals. In contrast, zebrafish are capable of scar-free regeneration of tissues that have historically been challenging to treat in clinic due to their low regenerative potential in mammals, such as the spinal cord, heart, and joints^1,4,6,7^. Current research in regenerative biology is dominated by techniques investigating the epigenome and transcriptome, but these powerful methods give an incomplete picture of regenerative processes. Filling this gap in knowledge about proteomic regulation of regeneration is important as proteins are the main drivers of cellular physiology. PTMs can generate multiple proteoforms from a single protein to serve a diverse array of cellular functions. Here, we investigate the landscape of post-translational protein regulation in both a regenerative and fibrotic model of ligament healing in the same animal model. This approach allows us to directly compare the differences associated with the distinctive healing outcomes. We demonstrate a role for Lgmn in facilitating the engagement between old and new matrix through regulation of a larger protease network.

We find that macrophages home to the injury edge of the ligament bundle and engage in focal proteolysis through upregulation of many lysosomal factors such as Lgmn, cathepsins, and MMPs (Figure 8). This localized degradation of mature ligament matrix primes the fibers for integration into the new scaffold. In the absence of Lgmn, we show that ECM-remodeling is dysregulated due to impaired degradation of Fn1a/b in the injured ligament. Failure to reorganize the ligament matrix and integrate with new matrix may contribute to the subsequent mis-patterning of the neo-ligament. Previously, we showed that *mpeg1.1+* macrophages home to the ligament injury border and express *lgmn* and *fn1a*^7^. Here we show that Lgmn is required to process Ctsla into its double-chain form, which has been previously identified as the prevalent isoform present in macrophages^43,44^. Macrophages could establish an adhesive interface by depositing new Fn1a proximal to the degraded ECM fibers in the ligament. During zebrafish cardiac regeneration, *mmp14b+* macrophages at the wound border are important for cardiomyocyte invasion into the injured tissue and expression of regenerative ECM components *col12a1a, fn1a,* and *fn1b*^2^. In the context of ligament injury, this Fn1a bridge could serve as a pathway for ligamentocytes to emerge out of the ligament and as a route for immune cells to infiltrate the injured ligament and secrete proteases.

**Figure 8.**
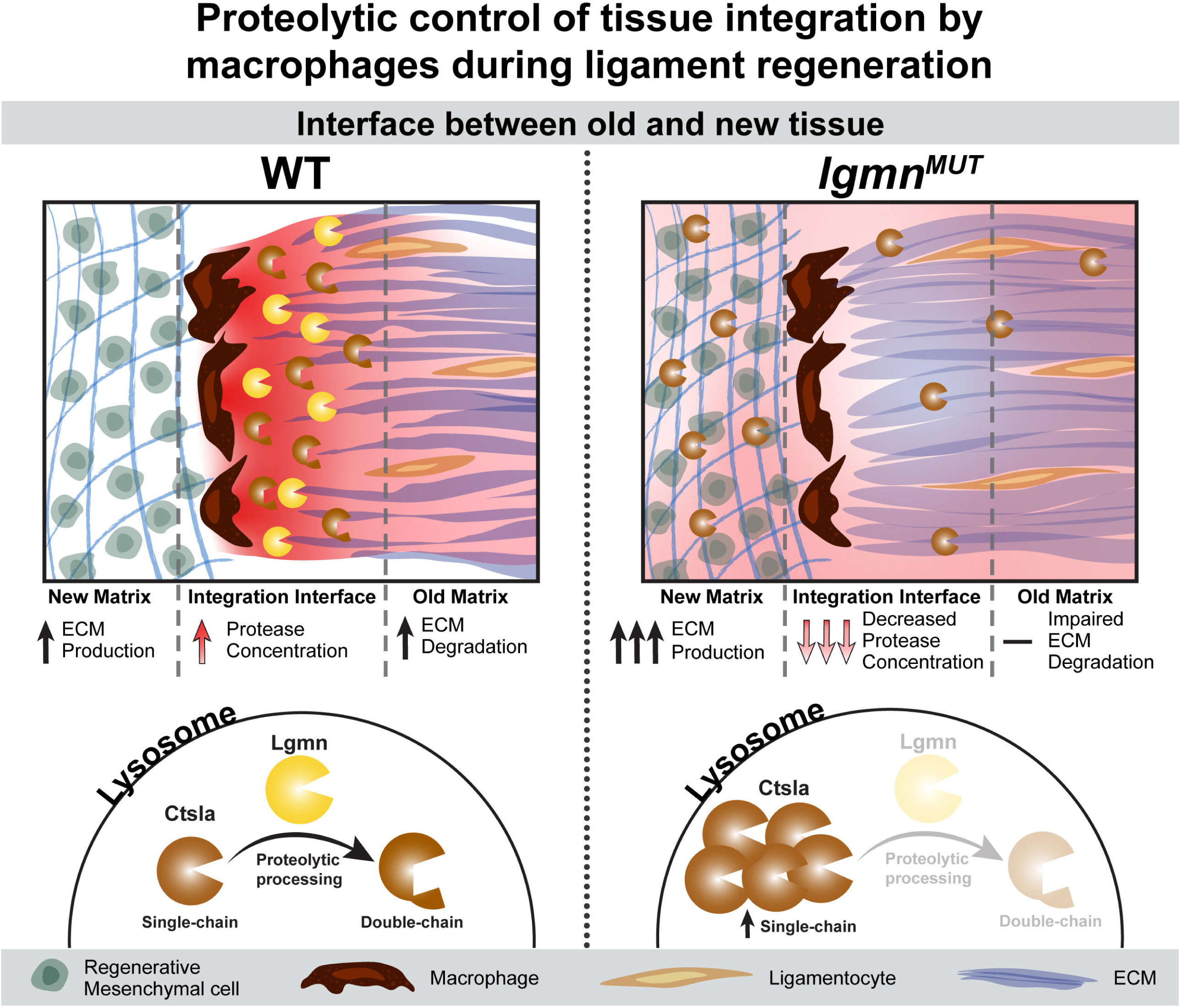
Model of proteolytic control of tissue integration by macrophages during ligament regeneration. Graphical model showing macrophage-mediated proteolytic processing of Ctsla (dark orange symbol) by Lgmn (yellow symbol) at the ligament injury edge for integration into the newly synthesized matrix. Red gradient indicates protease activity.

Together, these data suggest that macrophages are cellular architects of tissue integration through regulation of proteins involved in post-translational modifications. Wound healing requires a considerable amount of specialized ECM remodeling to occur in a constrained timeframe. As the ECM fluctuates in both composition and structure, it must continually support cellular processes such as migration and proliferation. Among these changes are increased expression of *fn1a* and *fn1b*^6,7^. Here, we show different expression profiles for the two paralogs associated with tissue organization. Furthermore, *fn1b* contains an additional FN3 domain, which may confer different structural properties and support different biological processes. Structural variation of fibronectin, including alternative splicing of mammalian fibronectin, alters its physical properties to support different cellular processes such as migration and proliferation during wound healing^45,46^. Proteoforms of fibronectin splice variants containing extra type FN3 domains have been shown to be specifically expressed during development, wound healing, and cancer^47,48^. This is due to a higher capacity for compaction and stretched conformations, thereby influencing cell-adhesion and fibril assembly^45,48^. Organization of fibronectin fibrils is important for tissue structure since the fibronectin scaffold is used as a guide for the deposition and assembly of other structural ECM proteins such as collagen^40,41^. In this study, we demonstrate that Lgmn is involved in establishing tissue organization through disassembly of Fn1a/b in the ligament matrix. Structural ECM proteins such as collagen, elastin, and fibronectin undergo critical PTMs that impart specific structural and mechanical properties to the matrix. Collagens undergo a series of hydroxylation and oxidation modifications required for crosslinking and maturation^49^. These modifications form degradation-resistant collagen that accumulate in murine hearts but not in zebrafish hearts, allowing the initial fibrotic scar to be reabsorbed^50^. Furthermore, enrichment of small-leucine rich proteoglycans modulate the mechanical properties of the injury matrix and has been associated with rodent and human spinal cord injuries but not in zebrafish^4^. Interestingly, *lgmn^MUT^* 3 dplt samples did not have an enrichment for factors involved in collagen lysine hydroxylation (P3h2, P3h3, and Plod1a) that were enriched in WT 3 dplt when compared to uninjured. Potentially, collagen in *lgmn^MUT^*may experience lysine under-hydroxylation, causing altered collagen cross-linking and altered mechanical properties of the tissue. Further investigation into the mechanical properties of *lgmn^MUT^* matrix during regeneration is required to fully understand the downstream consequence of altered post-translational modifications of collagens.

LC/MS proteome profiling uncovered a global decrease in protease levels in *lgmn^MUT^,* which surprisingly did not impair global protease activity. Although global protease abundance was decreased in *lgmn^MUT^*, this was not reflected in a change in functional activity because elevated levels of Ctsla compensated for Lgmn deficiency. While sufficient to support tissue development, we observed that compensation for lost Lgmn through upregulation of another lysosomal cysteine protease, Ctsla, is not sufficient for proper ECM remodeling required for integration in regeneration. This insufficiency in regeneration may be due to the absence of DC-Ctsla. This shift towards SC-Ctsla as the dominant species could influence stability, enzymatic potency, and subcellular localization, therefore altering rates of proteolytic processing and changes in substrate repertoire^51^. Unlike promiscuous proteases involved in non-specific cleavage, Lgmn is highly discerning with a narrow specificity for cleaving after asparagine residues in the P1 position and occasionally aspartic acid. In contrast, cathepsin L has a broader substrate profile, driven by a preference for aromatic and aliphatic residues in P2^32^. The irreversible nature of proteolysis requires stringent regulation of protease expression, activation, and localization to avoid detrimental consequences triggered by non-specific protein degradation. The restricted nature of Lgmn-mediated proteolysis and its high level of conservation in protein structure suggests that it has evolved a specialized role in regulating cellular physiology. This is evident in Lgmn and Ctsla localization at the injury border to remodel fibronectin fibrils and the role of Lgmn in processing cathepsin L, a potent collagenase and elastase^52–54^. Moreover, *lgmn^MUT^*exhibit a complete loss of multiple proteases (Cela1.3, C6ast3, Prss59.1) in response to injury, therefore further altering the composition of processed substrates. Cela1.3 (CELA1 in human) is a chymotrypsin-like elastase, C6ast3 (ASTL in human) is a metalloendopeptidase, and Prss59.1 (Prss1 in human)^55^ are all predicted to be involved in ECM degradation. Transcripts for some of these proteases were not detected in our scRNAseq, which can be explained by two possibilities: i) these proteins came from cells that were not captured in the sequencing due to technical limitations of FACS-mediated isolation or ii) their transcripts have degraded while the proteins have been placed in longer-term intracellular storage such as in secretory granules found in neutrophils and macrophages^56^.

This work highlights the importance of post-translational regulators as critical factors required to support regeneration. We show a requirement for Lgmn-mediated proteolytic regulation during early regeneration to facilitate proper ECM remodeling. Expanding research into regeneration-specific post-translational modifications is essential to better identify ways to better modulate human healing towards regeneration. Functional proteomic research is required to definitively state which proteins are present and PTM analysis to uncover the functional state of pro-regenerative proteins. Regeneration requires an orchestrated series of complex events, and here we show how Lgmn is a key specialized tool that controls an interconnected protease network at the wound boundary.

## Resource Availability

### Lead Contact

Requests for further information and resources should be directed to and will be fulfilled by the lead contact, Joanna Smeeton.

### Materials Availability

There were no new materials generated in this study.

### Data and Code Availability

• Mass spectrometry raw data files are deposited in MassIVE proteomics repository at https://massive.ucsd.edu/ and ProteomeXchange.org under dataset MSV000102222.

• Raw and processed scRNAseq data are deposited in GEO and can be retrieved using the accession number GSE335681.

• Raw western blots are deposited on Mendeley Data and can be accessed with DOI: 10.17632/8xkzs826vs.1

• Any additional information required to reanalyze the data reported in this paper is available from the lead contact upon request.

## Acknowledgements

We thank Joshua Barber for fish care. For scRNAseq experiments, we thank Michael Kissner at the CSCI Flow Core, Erin Bush, Simoni Tiano, and Joseph Mullen at the CUIMC Single Cell Analysis Core funded in part through the NIH/NCI Cancer Center Support Grant P30CA013696. This publication used the Columbia Genomics and High Throughput Screening Shared Resource supported by the National Center for Advancing Translational Sciences, National Institutes of Health, through Grant Number UL1TR001873. The content is solely the responsibility of the authors and does not necessarily represent the official views of the NIH. For LC/MS proteomics, we thank Michael Ford of MSBioworks. The authors’ work was supported by the National Institutes of Dental and Craniofacial Research (NIH R00DE027218 to J.S., F31DE033220 to J.M., F31DE031970 to T.A), T32 training programs NIGMS GM007088 (T.A. and J.M.) and NIAMS Integrated Musculoskeletal Training Program (IMTP) 5T32AR080744 (S.A.), Barnard SRI Funding to H.L., National Science Foundation NSF GRFP to M.B., Dr. Ines Mandl Connective Tissue Fellowship to K.S., and Columbia Stem Cell Initiative Seed Fund Award to J.S.

## Author Contributions

J.S. and J.M. conceived the study. J.S., J.M., and H.L. developed methodology. J.M., H.L., S.A., S.L., D.S., T.A., M.B., E.W., and K.S., performed investigation. J.M., H.L., and J.S. designed the visualization. J.S. administered and supervised the project. J.M. and J.S. wrote the original draft. J.M., M.B., E.W., K.S., and J.S. revised and edited the manuscript.

## Methods and Materials

### Zebrafish lines

The Institutional Animal Care and Use Committees of Columbia University approved all zebrafish experiments. Published Zebrafish lines used in this study include, *AB, Tübingen, lgmn* (sa22350)^57^, *Tg(scxa:mCherry)*^58^. Embryos were raised in a methylene blue salt solution at 28.5°C. Juvenile and adult fish were housed in groups of 15-30 individuals. All surgical injuries were performed in size-matched 3-12 months post fertilization (mpf) adult zebrafish using standard body length (SL) measurements. *lgmn^sa223^*^50^ mutant allele contains a G>T point mutation that encodes for a premature stop codon in exon 2. The truncated protein would encode for a 52 aa peptide. Homozygous *lgmn* mutants will be referred to as *lgmn^MUT^.* Zebrafish were genotyped with the method listed in Anderson and Mo et al., 2023^7^.

### Adult ligament transection surgery

The interopercular-mandibular (IOM) ligament transection was performed as described in Smeeton et al., 2022^59^. Briefly, adult zebrafish were anesthetized with Tricaine MS-222 and the IOM ligament was transected with a single cut using 3-mm Vannas spring scissors (Fine Science Tools, cat. 1500000). Fish were revived in clean system water and received post-operative health checks for the first three days after being housed on system.

### Histology, RNAscope, and immunohistochemistry

Adult zebrafish samples were fixed in 4% PFA at 4°C overnight, washed 2x30 min in 1xPBS and heads were isolated for embedding. Samples were decalcified in 0.5 M EDTA solution for at least 10 days rocking at RT. For paraffin embedding, the tissue was dehydrated in a series of EtOH dilutions, then the water was replaced with a Xylene substitutes Hemo-De (Electron Microscopy Sciences, cat. 23412-01) or ProPar Clearent (Rankin, cat. ANA-510) in a series of EtOH dilutions. Samples were incubated in a 1:1 Xylene Substitute:Paraffin solution at 65°C for an hour before incubating in 100% paraffin overnight. We used the Thermo HM255S automatic microtome to collect 5 μm sections.

For immunofluorescence (IF) staining, sections were deparaffinized, rehydrated through an ethanol series, and treated with sodium citrate (pH 6.0) buffer for 25 min at 90°C for antigen retrieval. Samples were blocked with ImmunoStain blocker solution (GeneTex, cat. GTX73323) for 1 hr then incubated overnight at 4°C in primary antibody diluted with blocker solution, see below for antibody information (Lgmn 1:250) (Lamp1 1:500) (Ctsl 1:200) (Fn1a/b 1:250). Day 2 of IF involved slides being washed in 1xPBST, incubated with secondary antibody ([1:500, Invitrogen Alexa Fluor] or [1:1000, Invitrogen Alexa Fluor Plus]) for 1 hr at RT, washed, and mounted with Fluoromount-G. Multiplex of tyramide signal amplification (tsa) and IF were done to co-stain using two antibodies raised in the same host. The Aluora Spatial Amplification HRP kit (Goat-anti-Rabbit, cat. A40001341) was used for tsa and was performed as per manufacturer’s instructions using. Briefly, slides were deparaffinized, rehydrated through an ethanol series, and treated with sodium citrate buffer (pH 6.0) for 25 min at 90°C in a steamer. Next, slides were washed with 1xPBS, treated with 3% hydrogen peroxide for 15 min, washed in 1xPBS, and blocked with the provided blocking buffer for 1 hr at RT before incubating overnight in diluted primary antibody at 4°C (Lgmn 1:2500) (Ctsl 1:2000). The next day, samples were washed in 1xPBS and incubated in poly HRP-conjugated anti-rabbit for 1 hr at RT, washed in 1xPBS, treated with Aluoradye for 8min at RT, washed in 1xPBS, and antibodies were stripped off with a 15 min treatment in sodium citrate buffer (pH 6.0) in at steamer at 90°C before continuing with IF protocol described above. Single-molecule fluorescence *in situ* hybridization was performed according to manufacturer guidelines using RNAScope Multiplex Fluorescent Reagent Kit v2 Assay (Bio-Techne). Slides were treated with 1x Target Retrieval Reagent for 4 min in a 90°C steamer. Opal 520, 570, and 690 fluorophores were used to visualize expression (1:1,000; Akoya Biosciences, cat. #FP1487001KT, #FP1488001KT, and #FP1497001KT). Slides were counterstained and mounted with Fluoromount-G + DAPI. RNAscope probes used in this study include: (Dr - lgmn (1003381-C2)), (Dr - mpeg1.1 (536171-C3)), (Dr - ctsla (1867391-C1)), (Dr - fn1a (1097911-C1)), (Dr - fn1b (840291-C3)).

### Imaging

Fluorescence imaging was preformed using a Leica SP8 confocal microscope with LASX software (v 3.5.7.23225). Fluorescent images were generated through maximum projection of a Z-stack, unless stated otherwise. Live imaging of transgenic line was performed using a Leica M165FC stereo microscope with Leica K5 sCMOS camera and LASX software (v 3.7.4.23463).

### Antibodies

Lgmn antibody (Cell Signaling Technologies, cat. 93627) [1:1000 for WB] [1:250 for IF] and [1:2500 for tsa]

Fibronectin 1a/b antibody (Millipore Sigma, cat. F3648) [1:1000 for WB] and [1:250 for IF]

Lamp1 antibody (Abcam, cat. ab24170) [1:500 dilution for IF]

Ctsl antibody (Genetex, cat. GTX128324) [1:5000 for WB], [1:200 for IF], and [1:2000 for tsa].

### Crystal structural alignment

Pymol was used to model the crystal structure of human proLGMN (pdb_00004fgu) determined from X-ray diffraction 3.9 Å^60^ and zebrafish proLgmn structure was predicted using Alphafold DB (AF-B8JHW0-F1-model_v4)^61^. RMSD value was determined through pairwise alignment in PYMOL (v 3.1.0).

### Lysate collection and western blot

Zebrafish jaw joints were resected and placed in ice-cold RIPA buffer containing Halt Protease and Phosphatase Inhibitor Cocktail and EDTA (Thermo, cat. 78440), manually homogenized with a micro-pestle, and centrifuged at 20,000 g for 20 min at 4°C. Supernatant was collected and samples were prepared in 1x BOLT LDS loading buffer and 1x BOLT reducing agent and boiled at 95°C for 5 min. 20μg of protein per sample was loaded into a Bolt Bis-Tris 4-12% gel (Invitrogen, NW04125BOX) and ran in 1x MOPS SDS running buffer (Invitrogen, cat. B0001) at 120V for 1.5 hrs. Proteins were wet transferred onto a methanol activated PVDF membrane with transfer buffer containing antioxidant supplement and 20% methanol at 4°C for 1.5 hrs at 90V. Total protein was stained by reactivating the membrane in 100% methanol and incubating in a 0.1%(w/v) PonceauS in 5% acetic acid for 5 min rocking. Ponceau-stained membrane was rinsed 3x in MilliQ and imaged with the iBright CL1500 imaging system. The membrane was blocked for 1 hr rocking at RT in 5% BSA in 1xTBST(0.1% Tween-20), washed 3x5 min in 1xTBST, and incubated overnight at 4°C rocking in primary antibody diluted in 5% BSA in 1xTBST. The membrane was washed 3x5 min in 1xTBST rocking at RT and then incubated for 1 hr at RT rocking in dilution of secondary antibody (Cell Signaling Anti-rabbit IgG, HRP-linked Antibody #7074) at a 1:2000 dilution in 5%BSA in 1xTBST. Followed by 5x5 min washes in 1xTBST before applying the SuperSignal West Dura Extended Duration Substrate solution (Thermo, cat. PI37071) or SuperSignal West Pico PLUS substrate (Thermo, cat. PI34580) and imaged for chemiluminescence on the iBright imaging system. Membrane was then stripped by incubating with 15mL of Restore Stripping Buffer (Thermo, cat. 21059) at 37°C rocking, washed 3x with 1xTBST and re-probed from blocking step as described above.

### Single-cell RNA sequencing

Ligament transection (n=16 joints per 1 dplt, 3 dplt, and 7 dplt time points) was performed on WT and *lgmn* mutant fish from 3-9 mpf. For sample collection, the whole jaw joint was resected from injured fish and uninjured samples (n=16 joints per genotype) and washed with ice-cold Ringer’s Solution in low-bind microcentrifuge tubes. Following resection, samples were mechanically dissociated with forceps in 28°C protease solution using collagenase D and collagenase P followed by nutation and agitation with a P1000 pipette every 5 min for 1 hr-1.5 hr until cells were in suspension. Live/dead cells were FACS sorted for using Draq7 (1:1000) viability dye on the Becton Dickinson FACSaria cell sorters. Gates selected live singlet cells and cells were collected in 100μl of collection media (2% FBS in 1xPBS). To determine cell concentration of sorted live cells, a portion of the samples was counted on the NovaCyte Quanteon using a mix of propidium iodine and Draq5 labels. The remaining sample was used to create single-cell RNA Sequencing libraries using the Chromium Single Cell 3’ Library and Gel Bead Kit (10X Genomics), according to the Chromium Next GEM Single Cell 3’ v3.1 protocol manufacturer’s recommendations. GEMs were generated in house by loading ∼15,000-16,000 cells for each sample using the 10x Chromium Controller, and the Columbia Single Cell Core conducted the downstream library preparation and processing according to NovaSeq manufacturer’s recommendation. Reads were aligned to the GRCz12tu genome assembly as reference using Cell Ranger-6.1.2.

Analysis of scRNAseq libraries were performed using R (v 4.5.1) in RStudio with Seurat (v 5.3.1)^34^. Seurat objects for each sample were made and merged into one object. We filtered for cells that had between 500 and 5000 features, less than 45,000 unique features, and less than 7.5% mitochondrial DNA to remove low-quality cells. Data was normalized before using PCA analysis to compute 80 principal components. Elbow plot used the PCA reduction to determine the dimensionality of the data. Principal components 1-50 were used for K-nearest-neighbor graph construction (KNN). Low-resolution (r = 0.9) clustering was used to determine broad cell types. Clusters and gene expression were graphed using UMAP-non-linear dimensional reduction to preserve global variability. Two individual objects were made using a subset of skeletal clusters and immune clusters, respectively. Objects were then subjected to the same preprocessing described above. Further sub clustering of those objects was conducted (skeletal subset, resolution = 0.95) (immune subset, resolution = 0.7) before adding back into the parent object. The DESeq2 (v 1.50.2) package in R was used for differential gene expression analysis.

### Liquid chromatography-tandem mass spectrometry (LC-MS/MS) proteomics

For label free LC-MS/MS, 3 dplt injury samples were collected by resecting 10 whole joints for each sample and uninjured samples were collected by resecting 15 contralateral uninjured joints that were then flash frozen in low protein binding microcentrifuge tubes. Three samples for each condition were sent to MSBioworks (Ann Arbor, Michigan, https://www.msbioworks.com/) for tissue processing and mass-spectrometry. Tissue was lysed in modified RIPA buffer (50mM Tris HCl, pH 8.0, 150mM NaCl, 2.0% SDS, 0.1% TX100, 1X Roche Complete Protease Inhibitor) using 1.4mm stainless steel beads in a Next Advance Bullet Blender, 2 cycles x 3 min each.

Samples were heated to 60°C for 30 min and clarified by centrifugation at 13.2K rpm. Protein was TCA precipitated overnight at -20°C, the pellet washed, and resuspended in 8M urea, 50mM Tris HCl, pH 8.0, 1X Roche Complete Protease Inhibitor. Protein quantitation was performed using Qubit. 25μg of each sample was digested overnight with trypsin. Briefly, samples were reduced for 1 hr at RT in 12mM DTT followed by alkylation for 1 hr at RT in 15mM iodoacetamide. Trypsin was added at an enzyme:substrate ratio of 1:20. Each sample was acidified to 0.3% TFA and subjected to SPE using Waters μHLB. 500ng per sample was analyzed by LC/MS with a ThermoFisher Vanquish Neo UPLC system interfaced to a ThermoFisher Orbitrap Astral. Peptides were loaded on a trapping column and eluted over a 150μm analytical column at 1.5μL/min; the column was heated to 55°C. A 30 min gradient was employed. The mass spectrometer was operated in data-independent mode. Sequentially, full scan MS data (240,000 FWHM resolution) from m/z 380-980 was followed by 300 x 2m/z precursor isolation windows; products were acquired in the Astral at 40,000 FWHM resolution. The maximum ion inject time (IIT) was set to 3.5ms for DIA; the NCE was set to 25. DIA data was analyzed using DIA-NN (v 1.9.2) to convert RAW files to QUANT, align based on retention times, search data using *in-silico* spectral library created from the FASTA file and iteratively using the spectral library created from RAW data, filter database (Uniprot Zebrafish) search results at 1% peptide and protein false discovery rate (FDR), calculate peak areas for detected peptides (Peptide Length = 7-30AA), and data normalization.

### LC/MS data processing and analysis

For LC/MS data processing, all statistical analysis was conducted on processed data after missing values were filtered out, intensity was log transformed, normalized, and imputed. When filtering out missing values, we used the following logic: for the identification of genotype-specific (WT vs *lgmn^MUT^)* and injury-specific changes in protein abundance (uninjured vs. injury), we included proteins where at least one experimental condition had 2/3 valid values. However, to include proteins undetectable in *lgmn^MUT^* such as Lgmn or injury samples, we included proteins that showed invalid values in all three replicates of at least one experimental condition. We used the DEP package (v 1.32.0)^62^ in R to log transform the intensity readings and normalize the data. To run differentially enriched protein (DEP) analysis, missing values were imputed from a manually defined left-shifted Gaussian distribution (shift = 1.8 and scale = 0.3) to account for values missing not at random. DEP package was used to run PCA and DEP analysis. Significant DEPs were determined (p-value ≤ 0.05 and LFC > 1 or <-1) and used to perform an unranked STRING (v 12.0) analysis to identify protein-networks and enriched pathways. Global protease abundance was determined by annotating proteases based on a reference list generated from UniProt using the keywords “((taxonomy_id:7955)) AND (keyword:KW-0645)” accessed on October 10^th^, 2025, and the reference table was deposited into MassIVE. R was used to annotate proteases identified by LC/MS and the global protease abundance was determined by taking the sum intensity values for all proteases and normalizing to the sum intensity value for all proteins for each sample. Proteins were classified as post-translational modifiers if they were in the list of proteases generated from uniport or filtered for if the name included the following words: kinase, phosphatase, SUMO, methyltransferase, demethylase, acetyltransferase, deacetylase, glycosyltransferase, peptidase, hydrolase, protease, ligase, hydroxylase, dioxygenase, ubiquitin, or deubiquitinase. Proteins were then manually checked and confirmed as post-translational modifiers.

### Multimodal non-negative matrix factorization (NMF) analysis

The list of 8,211 shared proteins/genes identified by LC/MS proteomics and scRNAseq was used to link the two modalities across dimensions for NMF analysis. The multiNMF function from the geneNMF package (v 0.9.2)^63^ in R was used to distinguish consensus modules across multiple k values (k=5-15) over a list of samples, WT and *lgmnMUT* uninjured, 1, 3, and 7 dplt. Gene programs were identified with the specificity weight set to 7, weight explained at 0.5, and 15 set to the number of target programs. Modules with low silhouette and specificity scores were excluded from analysis. Each cell identified by scRNAseq was then ranked for their highest scoring module for downstream analysis.

### Protease Activity Assay

Protein lysates were generated by collecting 6 whole joints in ice-cold 1x Pierce Luciferase Cell Lysis Buffer (ThermoFisher, cat. 16189) and pulverizing them with a micro pestle. After incubating on ice for 20 min, they were centrifuged for 20 min at 20,000 g at 4°C. Supernatant was collected and each experimental replicate consisted of 25μL of protein lysate with 75μL of either 1x Digestion Buffer (pH 7.4) or 20mM Sodium Citrate Buffer (pH 5.0). Each biological sample was run with 2 technical replicates. Protease activity was measured using the EnzChek Protease Assay, red fluorescence (ThermoFisher cat. E6639) in a black-walled glass bottom 96-well plate. Each sample was incubated with 100μL of the 10μg/mL BODIPY casein working solution diluted in either digestion buffer or sodium citrate buffer overnight at 28°C rocking. Plates were read using the Molecular Devices SpectraMax iD3 plate reader with the Em/Ex set to 589/619 for a fluorescence endpoint reading. The working solution background reading was subtracted from each experimental sample for analysis. Readings were then normalized to total protein input from each sample determined from BCA assay.

### Quantification and statistical analysis

Western blot quantification was determined by using measurements for total protein, and band intensity was measured using the iBright Analysis Software (v 5.4.0). Total protein for each sample was determined from the PonceauS staining and the normalization factor for each sample was determined by normalizing to the WT uninjured sample. Relative intensity levels were calculated by taking the Volume(Int.) measurement and dividing by the normalization factor.

Image analysis was preformed using ImageJ (v 2.16.0/1.54p) to measure the intensity distribution of Fn1a/b across the distance of the regenerate area by drawing a rectangular ROI around the regenerating ligament to capture the area adjacent to the RA to the tip of the IOP bone and used the plot profile function. The measurements representing the average pixel intensity in each column across the selection were then graphed in Prism and the simple linear regression of each genotype was calculated. RNAscope intensity was quantified using ImageJ. An ROI was drawn based on tissue morphology and intensity signal was measured as Raw Integrated Density (RawIntDen) and normalized to ROI area.

All statistical analysis was performed with Graphpad Prism 11. We performed 2-way ANOVA with Tukey’s multiple comparison test for western blot quantification, RNAscope image analysis, and protease activity assay. Statistics to determine global protease abundance from LC/MS was performed with a 2-way ANOVA with Uncorrected Fisher’s LSD.

## Supplemental Excel Table Titles and Legends

**Table S1. Counts table split by sample of scRNAseq.** Counts table of the number of cells contributed from each sample condition contributing to each cluster identified in scRNAseq data.

**Table S2. DEGs of WT vs lgmn mutant transcriptome for each timepoint.** Differentially expressed genes were determined by comparing the transcriptome of WT vs *lgmn* mutant samples of the same timepoint using DESeq2.

**Table S3. Enriched pathways of WT 1 dplt and *lgmn* mutant 1 dplt transcriptomes.** Enriched pathways identified by STRING-db (v 12.0) analysis of genes enriched in either WT 1 dplt or *lgmn* mutant 1 dplt transcriptomes when compared against each other.

**Table S4. Injury activated proteins in WT.** DEP analysis was used to determine injury activated proteins enriched in WT 3 dplt by comparing against WT uninjured.

**Table S5. Injury activated proteins in *lgmn* mutant.** DEP analysis was used to determine injury activated proteins enriched in *lgmn* mutant 3 dplt by comparing against *lgmn* mutant uninjured.

**Table S6. List of shared and genotype-specific injury activated proteins.** List of significantly enriched proteins (p-value <= 0.05, Log2FC >= 1 or <=-1) identified in each sample for the listed comparison. List of Shared Proteins (Column H) identifies the proteins present in both WT 3 dplt (Column C) and *lgmnMUT* 3 dplt (Column F) injury activated proteins when comparing against uninjured of the same genotype.

**Table S7. Enriched pathways of WT injury activated proteins.** Enriched pathways identified from STRING-db (v 12.0) analysis of proteins significantly enriched in WT 3 dplt when compared against WT uninjured.

**Table S8. Enriched pathways of *lgmn* mutant injury activated proteins.** Enriched pathways identified from STRING-db (v 12.0) analysis of proteins significantly enriched in *lgmn* mutant 3 dplt when compared against *lgmn* mutant uninjured.

**Table S9. Genotype-driven differences of the regenerate proteome.**Differences in proteomic composition of WT and *lgmn* mutant regenerates were identified using the DEP package in R to compare WT 3 dplt against *lgmn* mutant 3 dplt.

**Table S10. Enriched pathways of genotype-driven differences of regenerate proteome.** Enriched pathways identified from STRING-db (v 12.0) analysis of proteins significantly enriched (p-value <= 0.05, Log2FC >= 1 or <= -1) in either WT 3 dplt or *lgmn* mutant 3 dplt when compared against each other.

**Supplemental Figure 1. l*g*mnMUT upregulate ECM components by 1 dplt.**

a) UMAP of scRNAseq clusters present in each sample in UMAP.

b) Expression of cluster marker genes.

c) Enriched GO pathways identified by STRING analysis at 1dplt in WT when compared to *lgmn^MUT^*.

d) Enriched GO pathways identified by STRING analysis at 1dplt in *lgmn^MUT^* when compared to WT.

e) Expression of Tgfβ-associated DEGs at 1 dplt.

