## Supplementary figures and images for "Scarless regeneration requires post-translational proteolytic regulation via macrophage-derived Legumain"

### Supplemental Figure 1

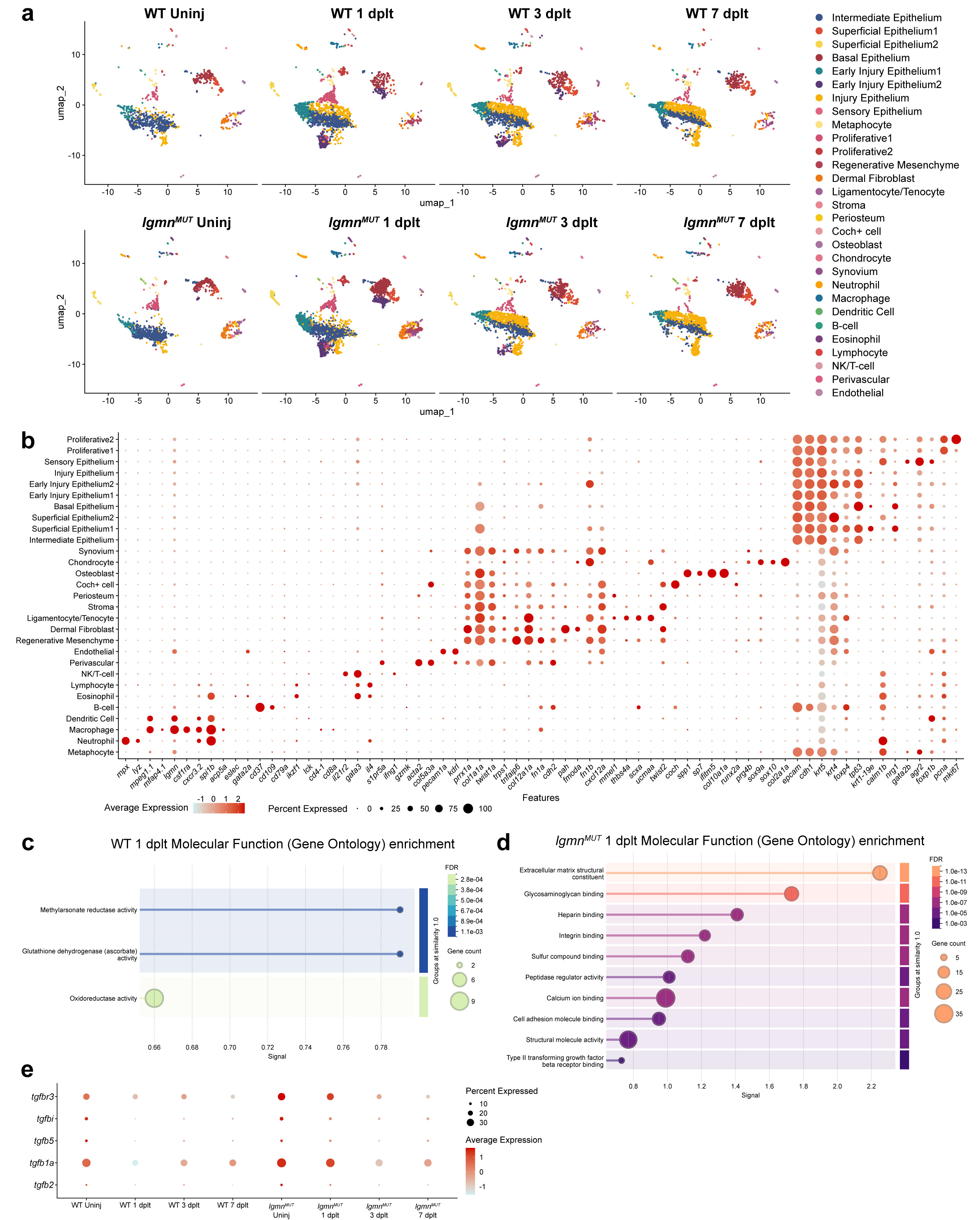
